# An algorithm to model the non-random connectome of cortical circuitry

**DOI:** 10.1101/2025.05.22.655546

**Authors:** Michael W. Reimann, Daniela Egas-Santander

## Abstract

Neuronal connectivity has been characterized at various scales and with respect to various structural aspects. In models of connectivity, it has so far remained difficult to match all of them at once, in particular the higher-order structure appears to be elusive. Here we introduce a new type of graph model that matches non-random structure characterized and described as relevant in biological neuronal networks. The structure emerges because the algorithm considers the need for axons to physically bridge the gap from soma to dendrites. If it targets one neuron, probabilities that it also targets other nearby neurons increase. We demonstrate that the algorithm can be successfully fit to complex, biologically relevant reference connectomes. Furthermore, we outline an intuitive expansion of the model from merely local to a combination of local and long-range connectivity. We provide a performant implementation that can be used to instantiate point neuron or morphologically-detailed network models at whole-cortex scale.

## 1 Introduction

The computational function of cortical circuitry is to a large degree implemented in the structure of its synaptic connectome. Yet, despite being studied for decades [1], much remains unknown about its structure. With respect to the rules governing its representation as a graph (the wiring diagram), we mostly know that simple algorithms do not capture its complexity [2, 3], although principles such a homophily [4] appear to provide better fits. Many investigations into this topic add complexity by introducing classes of neurons and formulating different rules for them [5, 6], i.e., they add complexity at a coarse-grained level from experimental observations. However, we also know that there is important structure on the per-node level, such as degree distributions [7], overexpression of reciprocal connectivity [8], of triad motifs [9], and larger motif classes, such as directed simplices [3, 10, 11].

In simulation-based neuroscience, models of neuronal networks are constructed and used to generate predictions of neural activity and the principles underlying it. This is often done in models that do not capture the known complexities of neuronal wiring due to the remaining uncertainties and the lack of availability of algorithms. Markram et al. [12], Reimann et al. [13, 14] used predicted connectivity based on detecting appositions between morphological reconstructions of axons and dendrites, which recreates an overexpression of reciprocal connections [2, 13], overexpression of triads [2], and partially recreates the overexpression of directed simplices [10, 15]. However, this algorithm requires an extensive library of morphological reconstructions and is in many contexts prohibitively costly to run.

Here, we introduce a new algorithm that is computationally cheap enough to wire a network of 10,000 nodes (i.e., neurons) in five seconds on a regular laptop computer. For a given number of nodes, it has only three parameters. We show that in parts of the parameter space, this class of graphs recreates the following features found systematically and reproducibly in biological neuronal networks: Long-tailed degree distributions, clustered connectivity as measured by a “common neighbor bias”, and overexpression of directed simplices. We propose that this arises in biological neuronal networks as a consequence of axons being required to physically reach dendrites when forming connections, which increases their probability to also innervate other nearby neurons. We demonstrate that this effect affects connectivity in an electron-microscopically measured connectome [16] in statistically highly significant ways and explain how our algorithm emulates the process.

Next, we introduce minor additions to the algorithm, determined by three additional parameters, that enable the customization of the coarse-grained and spatial structure of the resulting wiring diagram. These additions can be used to recreate the relative numbers of connections between classes of neurons (i.e., structural pathway strengths) and a directional bias for connections (e.g., more downwards than upwards connections). Both are aspects of connectivity that can be biologically measured and are arguably important. We then fit algorithm parameters against the wiring diagram of the connectome measured by the MICrONS initiative [16]. A large range of salient network measures of higher-order structure matched the reference, as well as the overall spatial structure of connectivity. We separately fit against (1) the subnetwork of excitatory neurons in layers 2/3 and (2) the subnetwork of excitatory neurons in layer 4/5 and found that their respective statistics can be recreated with largely identical parameter values, suggesting a universal wiring principle. To demonstrate the flexibility of the model, we also fit against the connectome of a morphologically-detailed network model [14] and recreate its statistics. Finally, we briefly outline an intuitive and logical extension of the algorithm to also capture the structure of inter-regional cortical connectivity. It is based on representing the locations of neurons not in cartesian space, but in a conceptual coordinate system based on inter-regional connection strengths.

## 2 Methods

### 2.1 Stochastic graph models

In this paper, we consider finite directed graphs without loops or double edges. Such a graph *G* is represented by a tuple (*V, E*), where *V* is the set of nodes and *E* ⊆ *V* × *V* is a set of directed edges, with no repeated entries (i.e., (*v, v*) ∈*/ E* for any *v* ∈ *V*).

We define a generative model of random graphs that depend on a single parameter *q*, called the *stochastic spread model on a base graph G* and we denote an instance of such a process by S*_q_*(*G*). Intuitively, a graph of this type is built from *G* by iteratively spreading from each node of *G* to a subset of its outgoing neighbors, which are selected at random. The size of the subset is also random with an expected value of *q* (Fig. 1A). The process then spreads further into the neighborhood of the newly selected nodes to a new set, and so on. Once a node has been a candidate to spread to once, it will be ignored in all future steps. As the size of the selected subset is random at each step, the empty set is a possible outcome, at that point the process stops. Additionally, the set of excluded nodes grows at each step, also forcing an eventual stop. In the output graph, an edge is placed from the starting node to all nodes reached by the spread. This process is then repeated for all nodes of *G*.

**Fig. 1:**
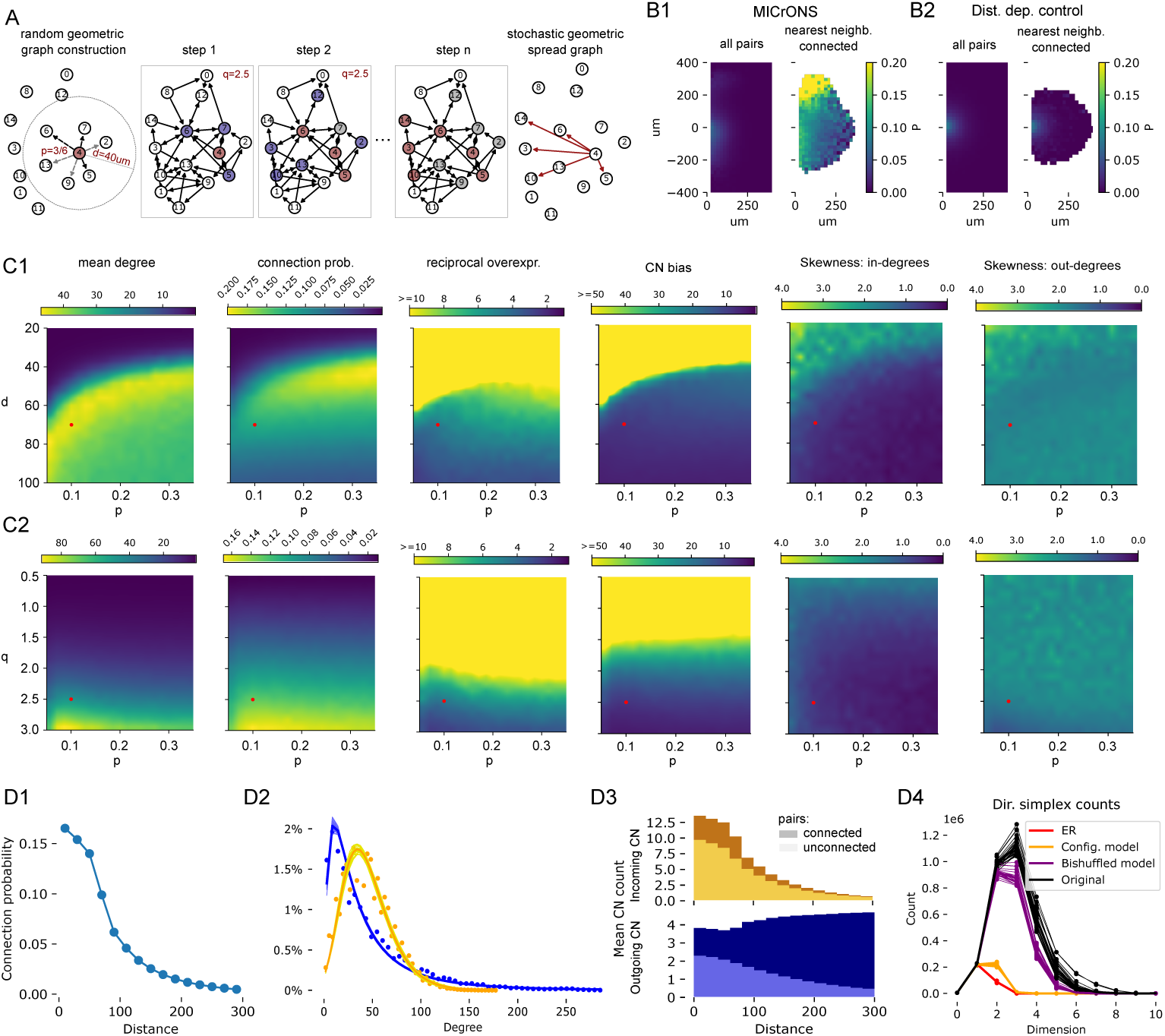
The stochastic geometric spread graph. A: Construction of a stochastic geometric spread graph on node 4. From left to right: A node is associated with a location and other nodes within distance *d* are detected (grey). A directed edge is placed to connect to them (black) with probability *p* (random geometric graph). Step 1: For a given node (red) the spread process selects its neighborhood (blue) as candidates. Nodes are selected from them randomly with an expected number of *q* nodes selected. Step 2: The neighborhood of the newly selected nodes (red) comprises the new set of candidates (blue), but nodes than were previous candidates but not selected (grey) are excluded. New nodes are selected from the new candidates. Step *n*: The process terminates when no new nodes have been selected. In the stochastic geometric spread graph, an edge exists from the starting node to all nodes reached by the process. B: Demonstration that this physicality of the axon affects connectivity. From left to right: Outgoing connection probability (20*µm* bins) in the connectome of the MICrONS initative for excitatory neurons in layers 4/5, and for horizontal (x-axis) and vertical (y-axis) offsets from the soma. Next to it, connection probability conditional on there being a connection from the neuron at 0, 0 to the nearest neuron in the spatial bin in question. Last two panels: Same, for a distance-dependent control connectome fit to the data. C and D: (caption continues on the next page). C1: Values of metrics characterizing overall connectivity and higher-order structure of SGSGs with different values of parameters *p* and *d*. For explanation of the parameters and metrics, see Methods. C2: Same, but for parameters *p* and *q*. Red dot indicates location in the parameter space of the instances analyzed further in D. D: (next page). D: More detailed analyses of parameter combination *p* = 0.1, *d* = 70*µm*, *q* = 2.5. D1: Distance-dependence of connection probability. D2: Distribution of in- (orange) and out- (blue) degrees. Dots indicate mean over 25 instances. Thin lines indicate lognormal fits for individual instances, thick line lognormal fit to pooled data. D3: Common neighbor bias: Number of common graph neighbors in SGSG instances for pairs at indicated distances. Light colored: for unconnected pairs, dark: For connected pairs. Top: incoming, bottom: outgoing common neighbors. Mean over 10 instances indicated. D4: Counts of directed simplex motifs in SGSG instances (black) and various controls fit to it. For details on the controls, see Methods. Each line one instance.

To formally describe this process, let N*_G_*(*v*) denote the outgoing neighbors of *v* ∈ **V**, i.e., all *w* ∈ **V** such that there is a directed edge in (*v, w*) ∈ **E**. For a subset of nodes *W* ⊆ *V*, we denote by N*_G_*(*W*) the all the outgoing neighbors of the nodes in *W* i.e., N*_G_*(*W*) = *_w_*_∈_*_W_* N*_G_*(*w*). Furthermore, we denote by N*_G,r_*(*v*) ⊆ N*_G_*(*v*) a random subset where each node is selected independently at random with probability *r*. If the base graph is understood from the context, we simply write N (*v*), N (*W*) and N*_r_*(*v*) for simplicity.

Now, for each node *v* of *G*, we will recursively define three subsets of nodes, which in step *i* are: the *candidate nodes C_i_*(*v*), the *selected nodes S_i_*(*v*) and the *tested nodes T_i_*(*v*). For *i* = 1 these are given by:

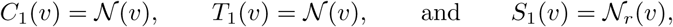

where 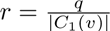. That is, *S*_1_(*v*) is a random subset of the outgoing neighbors of *v* of expected size *q*.

For *i >* 1 we define

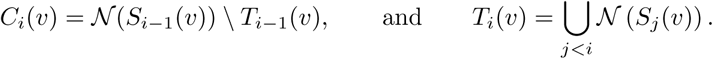

That is, the candidate nodes are the outgoing neighbors of the nodes selected in the previous step which are not nodes that have already been tested in any other step. Then, *S_i_*(*v*) ⊆ *C_i_*(*v*) is the subset where each node *u* ∈ *C_i_*(*v*) is selected independently at random with a probability proportional to the number of neighborhoods it appears in and such that the expected number of nodes selected is *q*. Thus, each node is selected with probability *r* ∗ *m_u_*, where *m_u_* is its multiplicity of the appearance of *u* across neighborhoods i.e.,

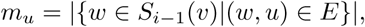

and

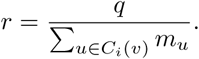

Finally, *S*(*v*) = *_i_ S_i_*(*v*) and S*_q_*(*G*) is a graph with nodes *V* which contains a directed edge from *v* to each node in *S*(*v*) for each *v* ∈ *V*.

Note that a stochastic spread graph on the empty graph is again the empty graph, while on the fully connected graph on *n* nodes is an Erdos-Renyi graph with overall connection probability 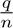. Moreover, this processes can be efficiently implemented based on sparse matrix multiplications.

Additionally, we also consider a type of *random geometric graphs* [17] on a set of points **P** in *n*-dimensional space, which we denote by *G_d,p_*(**P**). Such a graph has as nodes the set of points in **P** and directed edges added independently at random from *v* to *w* with probability *p* if the euclidean distance between *v* and *w* is smaller than *d*. We call a stochastic spread graph on such a random geometric graph, S*_q_*(*G_d,p_*(**P**)), a *stochastic geometric spread graph*, which we denote as SGSG.

### 2.2 Customizations of the models

We define a number of additional steps to the process that enable customization of the spatial and coarse-grained structure of the resulting SGSG graph. In the context of a connectome model for neuronal circuits, these biases can be used to match neuron type-based or spatial observations in biological connectomes.

#### 2.2.1 Biasing the spatial neighborhood selection

Above, we constructed a simple random geometric graph by placing connections to nodes in the geometric neighborhood of each vertex *v*, i.e., the set of nodes at euclidean distance smaller than *d*, which we denote by *H_d_*(*v*), independently at random with probability *p*. We introduce two types of bias into the process. Each bias defines a weight for every edge (*v, w*), which increases or decreases the probability that the pair is selected. These probabilities are then scaled by their corresponding weights and normalized such that the mean probability over all nodes in *H_d_*(*n*) remains equal to *p*.

**Per node biases** The simplest form of bias is to define for each node *v* a tuple of weights (*w_o_*(*v*)*, w_i_*(*v*)), which increase or decrease the relative out- and in-degree of the node *v*. To do so, we bias the probability of choosing each each edge (*v, u*) by the weight *w_o_*(*v*) · *w_i_*(*u*). While this bias has potentially different values for each node, in practice we used identical values for large groups of nodes, specifically for nodes representing neurons belonging to the same type.

**Orientation-based biases** This bias is determined by directional alignment with a base unit vector 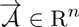, which we call the *orientation axis*. For two nodes *v* and *u*, corresponding to points **p***_v_,* **p***_u_* ∈ **P**, the probability of forming an edge (*v, u*) is maximally increased when the vector from **p***_v_* to **p***_u_* is parallel and aligned with 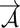; it is maximally decreased when the vector points in the opposite direction, and remains unbiased when it is orthogonal to 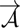.

More precisely, we choose a weight −1 ≤ *w* _A_ ≤ 1. Then for any *u* ∈ *H_d_*(*v*) let 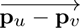 denote the vector from *v* to *u*. Then, the bias weight for (*v, u*) is:

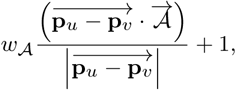

where · denotes the vector dot product. Note that the bias weight ranges from 1 + *w*_A_ to 1 − *w*_A_, depending the level of alignment with 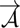.

#### 2.2.2 Reducing stochasticity of *S_i_*(*n*)

Above, we describe a process where *S_i_*_+1_(*v*) is generated from *S_i_*(*v*) by picking from outgoing neighbors in a random geometric graph independently at random such that the expected size of *S_i_*_+1_(*n*) is *q*. Here, we introduce an alternative method where instead exactly 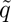 candidates are picked, where 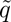 is *q* rounded to the nearest integer. This is similar to the two variants of an Erdos Reyni graph, one with a fixed number of edges and one with a fixed probability of having an edge between a pair of nodes [18, 19]. As there is no variability in the size of *S_i_*(*v*), we consider this a less stochastic version of the process. We employ the less stochastic version in the first *k* steps, i.e., for *S_i_* with *i < k*, then switch to the regular version for the remaining steps.

In the remainder of the manuscript we will use the asterisk to indicate graphs that have been generated with any of these modifications, i.e., S*_q_*(*G_d,p_*(**P**)) vs. S^∗^*_q_*(*G_d,p_*(**P**)), and SGSG vs. SGSG*.

### 2.3 Extension to long-range connectivity

We extend the SGSG to a proof-of-concept version that incorporates additional long-range connections. To achieve this, we generate two random geometric base graphs over the same set of nodes in 3-dimensional space: one with local connectivity constructed as before, and another where distances are defined by transformed coordinates of the original point could, that reflect the structure of long-range connectivity.

More precisely, from a set of points **P** ∈ R^3^, we generate another set of points **P**^∼^ ∈ R^3^ with transformed *y*-coordinates, while keeping the *x* and *z* coordinates fixed. The modified *y*-coordinates are given by 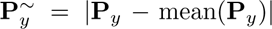, i.e., the distance from an *x, z*-plane at the center of the point cloud.

Then, we build two random geometric graphs, *G_d,p_*(**P**) and *G_d,p_*_·*m*_(**P**^∼^) on the same set of nodes, where *m* is a parameter that determines that balance between the local and long-range edges in the final result. We consider the graph given by the union of edges of *G_d,p_*(**P**) and *G_d,p_*_·*m*_(**P**^∼^) and denote it by 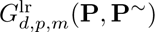. Finally, we build a stochastic spread graph on it, 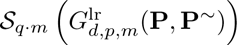. Note that other methods to determine **P**^∼^ and versions in which the point cloud is in *n* dimensional space, are possible and will change the overall and spatial structure of the long-range edges added.

### 2.4 Experimentally measured connectome

As a benchmark to compare algorithmically generated graphs against, we use the wiring diagram of the MICrONS connectome [16]. Specifically, we use the internal connectivity of release version 1181 of the data. Only connections between neurons that both had somata inside the volume were considered. As we are interested in the structure of the graph representation of connectivity only, we used the representation of the MICrONS data at cellular resolution of [15]. For classes of neurons in the data we used the table “aibs metamodel mtypes v661 v2” also provided by the MICrONS initiative [20]. Subnetworks of excitatory neurons in layers 4/5 and of excitatory neurons in layers 2/3 were separately analyzed. For access to that data, see Data and Code availability below.

### 2.5 Graph measurements used

- **Mean degree** refers to the mean number of incoming or outgoing edges across all nodes. Note that this value is the same regardless of using incoming or outgoing edges.
- **Connection probability within** 75*µm*: As nodes in our models are associated with spatial locations, we can calculate the probability that an edge exists in a given direction between a randomly picked pair within 75*µm*.
- **Reciprocal overexpression** is calculated as the ratio of the number of reciprocal (or bidirectional) edges in the graph over the number in an Erdos-Renyi graph with the same number of nodes and edges.
- **Common neighbor (CN) bias** is the ratio of the mean number of common neighbors of connected over unconnected pairs of nodes. Node *c* is a common neighbor of *v* and *u* if an edge exists from *v* to *c* and from *u* to *c*. If *u* and *v* are reciprocally connected, they are counted on the connected side twice, if they are unidirectionally connected they are counted once on both sides, etc.
- **Skewness**: We calculated the skewness of the distributions of in- and of out-degrees of nodes using the implementation in the scipy.stats package. This measures how much the distribution is long-tailed on the right side (positive values) or left side (negative values). Symmetrical distributions have a value of 0.
- **Directed simplices**: We calculated simplex motifs in the constructed graphs and their controls. A directed *n*-simplex, is a motif on *n* + 1 neurons which are all to all connected in feedforward fashion. That is, there is an ordering of the nodes 0, 1*,... n*, such that there is an edge (*i, j*) whenever *i < j* (see Fig. S1A.). Additional edges are allowed to exists.

### 2.6 Graph control models

To assess the significance of graph measurements used, we compared to the following controls that use the same nodes but randomized edges as a reference:

- **Erdos Renyi (ER)**: Edges are present with a uniform probability fitted to the reference. They are placed statistically independently.
- **Configuration model**: Additionally, the in- and out-degrees of all nodes are preserved.
- **Bishuffled model**: Most edges are identical, only bidirectional connections are shuffled as follows. For each bidirectional edge a single direction is randomly chosen. Then, in the resulting purely unidirectional graph, a randomly selected subset of edges is made bidirectional to match the number of bidirectional edges in the reference.

See Figure S1B for examples of these control models.

## 3 Results

### 3.1 Non-random properties of neighborhood spread graphs

We explore stochastic neighborhood spread graphs (SGSGs) as models for the local connectivity of cortical circuits. These are graphs where edges from a source node to target nodes are iteratively placed using a stochastic process that spreads along the edges of a second graph on the same nodes (Fig. 1A, see Methods). This is motivated by the idea that connectivity is implemented by axons and dendrites that have to physically cross the distance from pre- to post-synaptic neuron. Once we know that it has reached a certain point in space, the probability that it also reaches neighboring points is high. Similarly, once the stochastic wiring process has reached a given node it only has to spread along one additional edge to reach any of its neighbors. While intuitive to grasp, we have to demonstrate that this effect actually affects connectivity in a measurable way. To that end, we analyzed the connectome of layers 4 and 5 of the MICrONS connectome (see Methods). We predicted that connection probabilities for a pair of neurons (*A* and *B*) should be higher than average if we know that a connection from *A* to the nearest neighbor of *B* exists. That is, the existence of one edge is statistically dependent on another edge. We confirmed this to drastically affect the MICrONS data, increasing connection probabilities from near zero to over 20% even at distances around 300*µm* (Fig. 1B1). In a fitted, stochastic, distance-dependent control model connections are formed statistically independently, hence the effect is not present (Fig. 1B2). A similar idea has been independently explored by Piazza et al. [21] who provide compelling arguments that a connectome model must take these factors into account. But they stopped short of providing an actual wiring algorithm. A similar idea has been used by Reimann et al. [22] in their model of cortical long-range connectivity. Specifically, they used spread along an underlying graph to model which cortical regions are innervated together by individual projection neurons.

We begin by evaluating this family of graphs with respect to several features of connectivity that characterize the connectivity of cortical networks. To that end, we built instances of SGSGs while iterating the values of the three parameters over a wide range. Instances were built for 5000 nodes that were randomly uniformly distributed in a cube with a side length of 500*µm*, corresponding to a density of 40,000 per *mm*^3^. Biological cortical cell densities are on the same order of magnitude [12, 23]. We note that the density of the nodes used affects the structure of the SGSG graph in principle, but this can be compensated through the *d* parameter: If we scale each side of the cube and *d* by the same factor *x* and keep the number of points unchanged, we will generate the same SGSGs, but at eight times lower density. Hence, we do not consider the density a parameter to explore.

First, we considered measures of the density of connections: The mean node out-degree as a measure of the overall number of connections; the connection probability within 75*µm* to understand to what degree connections are found specifically between nearby nodes; and the overexpression of reciprocal connectivity (Fig. 1C). We found that different parameter combinations lead to very different levels of sparsity, indicating that a wide range of types of network can be recreated. Notably, while the mean degree and probability within 75*µm* were correlated, they were also independent to some degree. Finding e.g., the same mean degree with lower connection probability means that connections are re-distributed to above 75*µm*, it therefore shows that different types of distance-dependence can be implemented. Significant overexpression of reciprocity was found throughout the parameter space tested, but it was highest for extremely sparse networks.

Next, we considered measures of non-random higher-order structure that have been demonstrated to be relevant in biological neuronal networks [3, 7]: Skewness of degree distributions, as indication that the distribution is long-tailed, and the common neighbor connectivity bias. A common neighbor of a pair of nodes is a third node that is connected to both of them. We define the common neighbor bias as the mean number of common neighbors of connected pairs of nodes divided by the same measure for unconnected pairs of nodes. Its value increases the more the connectivity is clustered. While we found the highest values of CN bias and in-degree skewness for location in parameter space that resulted in extremely sparse networks, they were still significantly elevated over values for more random networks in the rest of the space. Skewness of out-degrees was relatively constant at values around 2.

We explored the parameter combination of *d* = 70*µm*, *p* = 0.1, *q* = 2.5 further, generating 25 SGSG instances. We confirm that its connectivity is distance-dependent, and that both in- and out-degrees are lognormally distributed as in biology (Fig. 1D1, D2). As expected from panel C, that distribution of out-degrees has a longer tail. Further, connected pairs have a larger number of common neighbors at all distances (Fig. 1D3), indicating clustered connectivity. Curiously, the number of efferent common neighbors was not distance-dependent for connected pairs, but for unconnected pairs. The increase in common neighbor count for connected pairs was larger for efferent than for afferent neighbors.

**Fig. 2:**
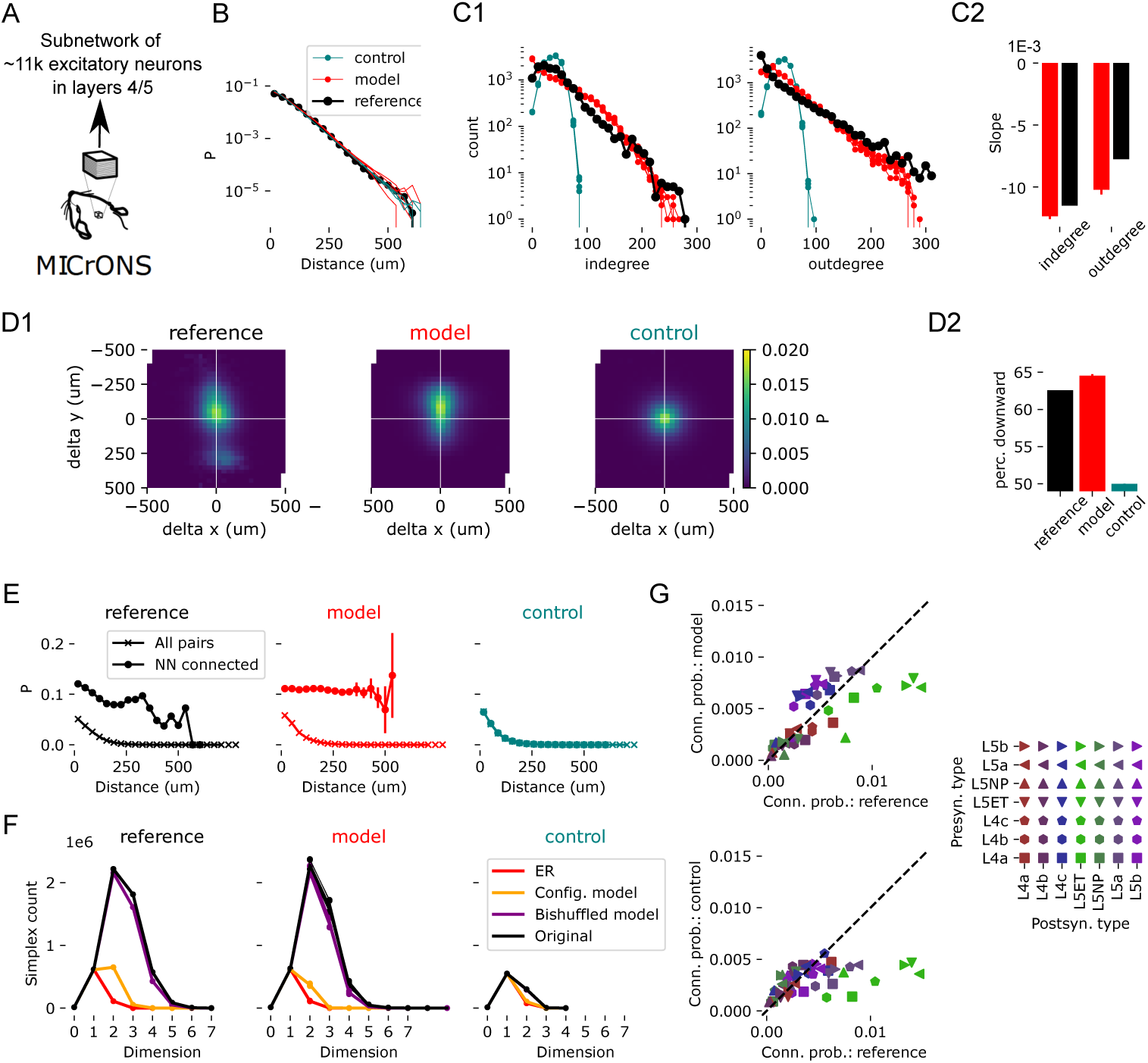
SGSG* fitted to the MICrONS connectome. A: As “reference” we used the network between excitatory neurons in layers 4 and 5 of the MICrONS data, our “model” is SGSG* instances fitted to the reference, the “control” is instances of a stochastic distance-dependent connectome fitted to the reference. B: Distance-dependence of connection probability in the reference (black), model (red, five instances) and control (teal, five instances). C1: distribution of in- (left) and out- (right) degrees in the connectomes. C2: Slopes of linear fits to the data in C1 for reference (black) and model (red). For the model, the bar indicates mean, error bar the standard deviation over five instances. D1: Connection probability in two dimensions for reference, model and control. The delta-y is roughly aligned with an axis orthogonal to layer boundaries, with negative values indicating downwards connections. D2: Fraction of connections going downwards in D1. Colors as in A. E: Analysis as in Fig. 1B, but considering only the overall distance. Data with x-marks indicates overall connection probability, data with circle-marks indicates connection probability conditional on the nearest neighbor being connected. F: Directed simplex counts as in Fig. 1D4. G: Pathway-specific connection probabilities, reference against model (top, pearsonr=0.7) and against control (bottom, pearsonr=0.45). Mean over five instances. Marker shape indicates pre-synaptic, color post-synaptic neuron types.

The analyses above measures non-random trends in the connectivity between three neurons at a time: One pair and their common neighbor. Larger motifs, called “directed simplices” have been shown to be overexpressed in virtually all biological connectomes and have implications for network function [3, 24, 25] (see also Methods for an explanation of these motifs). We found that in the SGSG instances simplices are highly overexpressed compared to strong controls (Fig. 1D4). Notably, their count is higher than even in a bishuffled control, that maintains most of the topology of the reference network and only randomizes the locations of bidirectional edges. This indicates that the locations of bidirectional edges are non-random with respect to the simplicial structure and has been previously shown to be the case in biological connectomes [3].

We conclude that SGSGs are a useful model for biological connectomes: First, the overall density of connections can be varied independently of the connectivity as close distances. Second, in a large portion of the parameter space, we find indication of higher-order structure that has been previously characterized in biological neuronal networks. Third, the values of metrics associated with this higher-order structure can be independently varied to some degree, enabling fitting to a reference connectome.

### 3.2 Fitting to the MICrONS data

We set out to use SGSG* as a model of specifically the connectivity of excitatory neurons in layers 4 and 5 of the MICrONS connectome (Fig. 2A; from here on: the “reference”). This means compared to the previous section, the following aspects changed. First, instead of randomly generated locations, we used the soma locations of the reference. Connection probability in the reference decreased more slowly along the y-axis (roughly orthogonal to layer boundaries), indicating a columnar organization of connectivity [26]. To match this, we simply divided the y-coordinates of neurons by a factor *f_y_* for the construction of the random geometric graph, thereby decreasing distances along that axis. Second, we used a number of additional customizations outlined in the Methods, that enabled us to match the spatial and more coarse-grained structure of the graph in addition to its topology. This added *per node biases*, *w_o_* and *w_i_*, that made connections from (*w_o_*) and to (*w_i_*) nodes more or less likely. These were respectively calculated based on the mean outgoing and incoming connection probabilities of the morphological type a neuron belonged to. Hence, this introduced parameters, but they were deterministically calculated and not subject to a fit. We also added an *orientation bias*, *w*_A_, of connectivity. We considered orientation along the y-axis that is roughly orthogonal to layer boundaries. We made downward connections (towards deeper layers) more likely, and those going upwards less likely. Finally, we introduced a number of steps with reduced stochasticity (*k* steps) in the generation of the stochastic spread graph S^∗^. During these initial steps, the stochastic process always spreads to a fixed number of new nodes, hence avoiding the presence of nodes with zero or very low out-degrees. The values of parameters were manually optimized and can be found in Table 1.

**Table 1:**
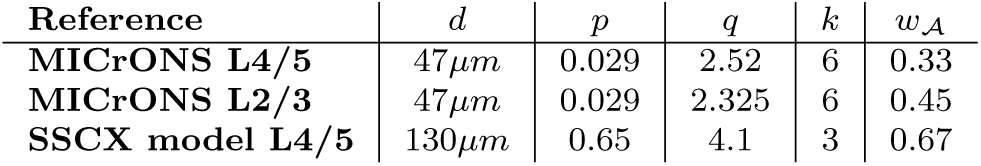
Manually fitted parameter values for different reference data sources.

We were able to obtain a great match to the reference for many relevant aspects of connectivity. Connectivity decreased with distance linearly in log-space (Fig. 2B), with a matching spatial bias making downwards-facing connections more likely (Fig. 2D). Degree counts were long-tailed unlike in a distance-dependent control (Fig. 2C). While our model had slightly steeper slopes, it qualitatively matched the observation that out-degrees were more long-tailed than in-degrees. We also found a match for increased connectivity when the spatial nearest neighbor is connected (Fig. 2E), the effect we explored in Fig. 1B. Counts of directed simplices have been previously used as a benchmark for biological neuronal networks, with significant overexpression found for organisms ranging from worm to rodent [3]. They have also been predicted to be functionally relevant [3, 24, 25]. In that regard, we also obtained a near-perfect match to the reference (Fig. 2F). Coarse-grained pathway strengths were reasonably well matched (Fig. 2G).

Next, we repeated these analysis, but instead used a different connectome as reference: layers 4/5 of the model of circuitry of rodent somatosensory region of Reimann et al. [14]. This model has been shown to have substantially non-random higher order structure [14]. This makes it a good benchmark to test the generalization capability of SGSGs, independent of its realism. We found that a very different set of parameters was required (Table 1), the original set led to twice sparser connectivity (not shown). After manual re-fitting of parameters we again obtained a good qualitative match to the reference (Fig. 3. A notable difference was that downwards facing connections had increasingly lower horizontal offsets with distance from the soma (Fig. 3D1, left), which was not reproduced. Curiously, in the one-dimensional connection probability, an upwards deviation from exponential decay was observed for distances above 500*µm*, which our model reproduced, albeit at an exaggerated level (Fig. 3B).

**Fig. 3:**
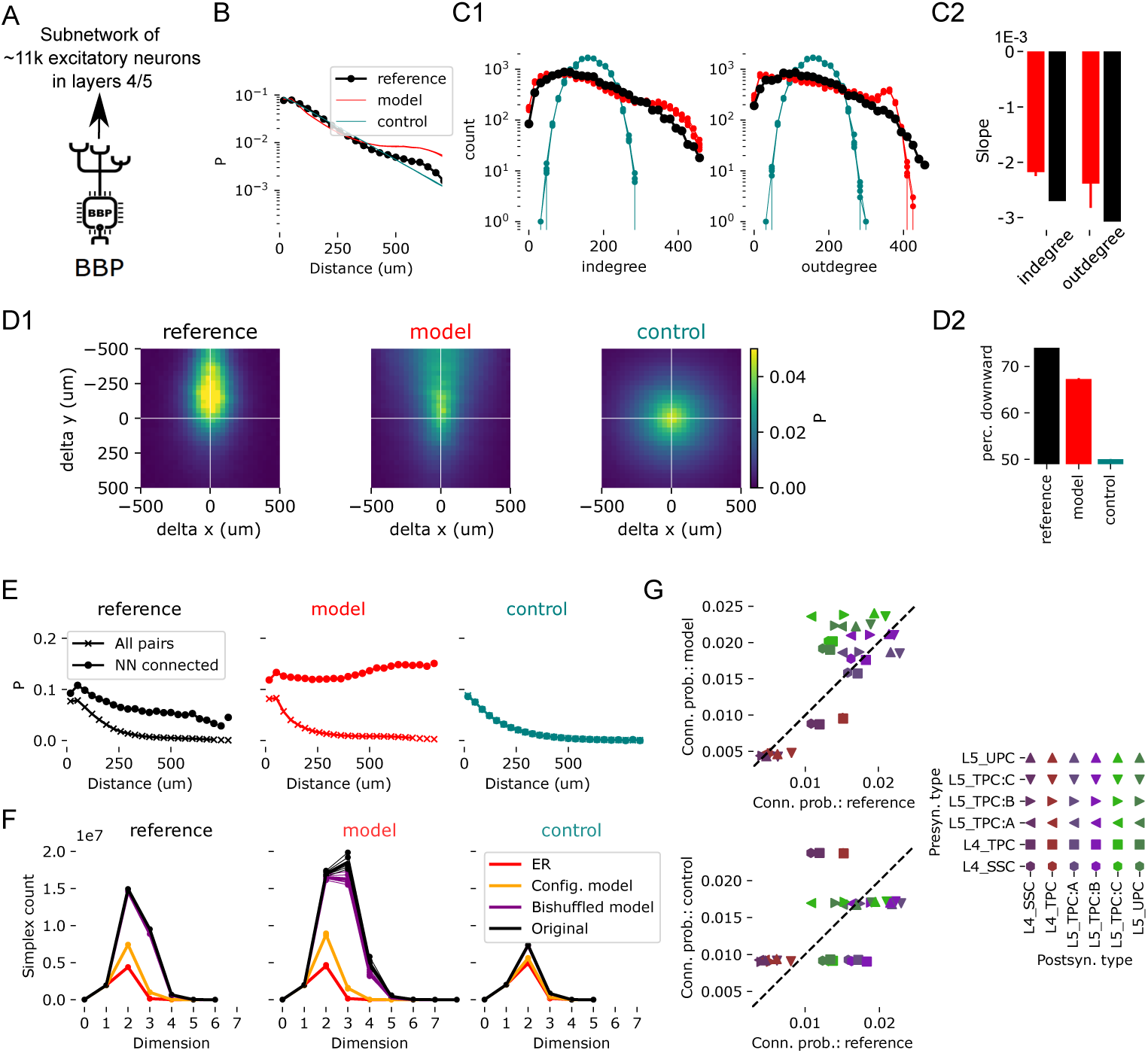
Comparison of a fitted SGSG* model to a morphologically-detailed model of connectivity in L4/5. Everything as in Fig. 2, but using connectivity in layers 4 and 5 the model of Reimann et al. [14] as reference. Parameters re-fit to the data. Pearsonr for the data in panel F: 0.80 for reference vs. model, 0.46 for reference vs. control.

We also repeated the analyses, for connectivity in layers 2 and 3 of the MICrONS data as reference (Fig. S2). Notably, we tested generalization, keeping three of the parameters (*p*, *d* and *k*) identical from the fit to MICrONS layers 4 and 5. *q* was re-fit to match the total number of edges in the new reference; *w*_A_ was re-fit to match the fraction of downwards facing edges; *f_y_* was re-fit to match the shape of two-dimensional connection probability indicated in Fig. 3D. The results matched the connectome overall qualitatively well, but not quite as well as for the previous reference. Most notably, in layers 2/3 the difference in the slope of in- and out-degree distributions was more pronounced, which the model captured incompletely (Fig. 3C2). We note that for different *p* and *d* a SGSG can generate these differences (see Fig. 1D2). Furthermore, the simplex counts in the reference followed a highly uncommon pattern, increasing from dimension four to six after decreasing from two to four, which the model did not reproduce (Fig. 3F). On the other hand, the model has a maximum dimension one above the reference and higher counts in dimensions two and three.

### 3.3 Extension to long-range connectivity

We believe that SGSGs can be extended to model both local and inter-regional (or long-range) connectivity at cellular resolution with some intuitive modifications. Briefly, we use the union of two random geometric graphs instead of a single. While the second one is built on the same nodes as the first, it uses different node locations (**P**^∼^) in its construction that encode the structure of inter-regional connectivity by proximity. Here, we explore a simple example (Fig. 4A): In an elongated point cloud we draw a region border simply as a dividing plane. One coordinate of **P**^∼^ is then the distance from that line. This will place nodes on opposite sides but at the same distance from the plane close to each other, leading to edges between them in the random geometric graph that could not be present when the regular locations (**P**) are used. This approach is based on topographical mappings of connections between primary and higher visual areas that follow such a rule in mouse [22, 27].

**Fig. 4:**
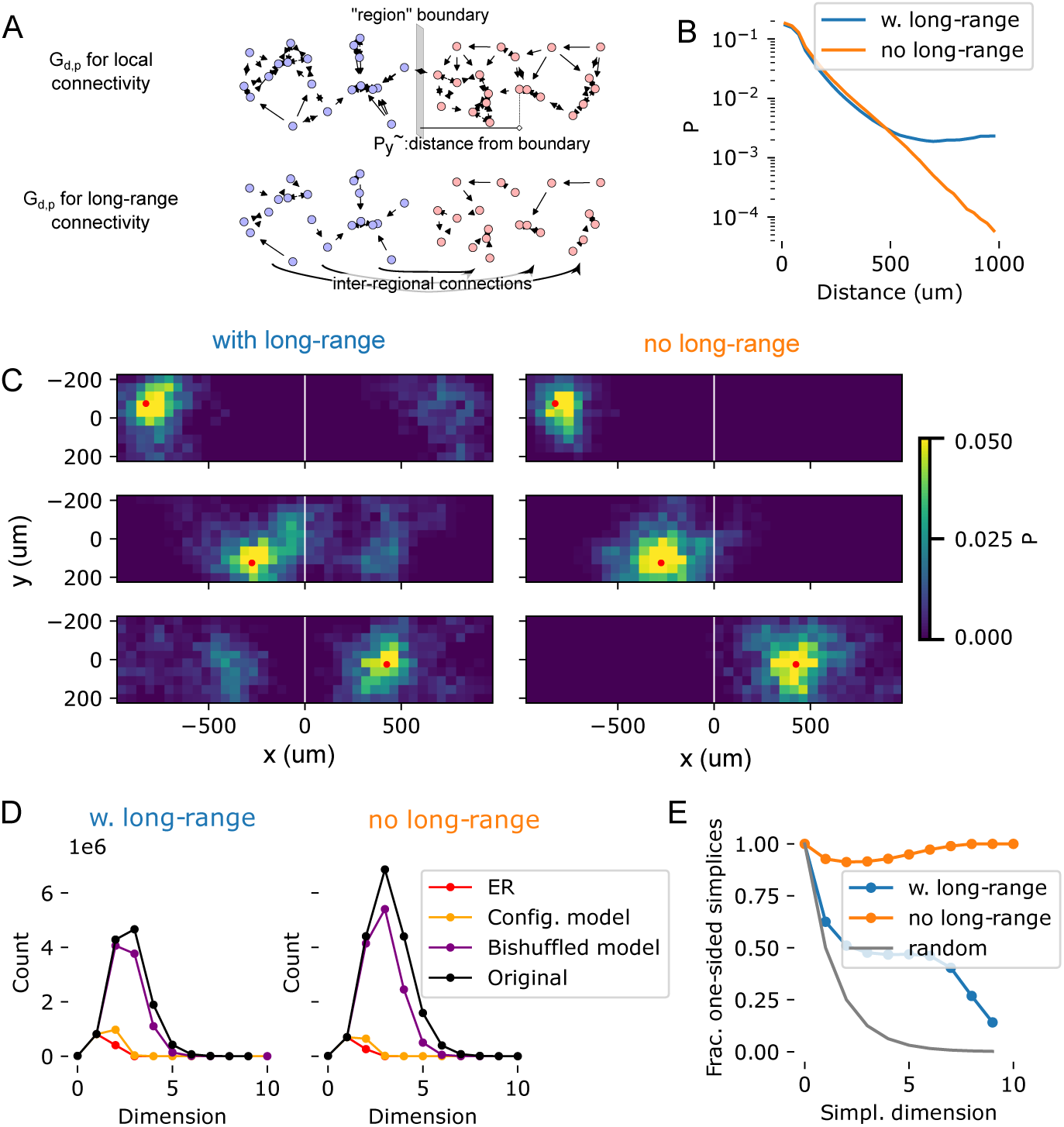
An extension to long-range connectivity. A: We extent the model by building two random geometric graphs, one for local and one for long-range connectivity, then using the union of their edges to build a stochastic spread graph. The long-range portion uses the same parameters, but “virtual” node locations instead of the real ones. Here, we use virtual locations based on the distance from a region border drawn in the middle of the points. B: Distance-dependent connection probabilities of the resulting SGSGs. Blue: As described in panel A; orange: built without long-range portion, as in Fig. 1. C: Two-dimensional connection probabilities of neurons in various spatial bins. Red dot indicates the location of a spatial bin, colors indicate the connection probabilities of nodes in that bin to all other bins. White line is the location of the region border. Left column: For three exemplary bins in a connectome model with long-range connectivity. Right column: For the same bins in a model without long-range connectivity. D: Simplex counts as in Fig. 2F with and without long-range connectivity in the model. E: Fraction of simplices that do not contain nodes on both sides of the region border. X-axis indicates the dimension of simplices considered. Blue and orange as in B. The grey line indicates results if the side of the border is shuffled for all nodes.

We built a stochastic spread graph on such a union of two graphs and contrast it with a stochastic spread graph on only the random geometric graph representing local connectivity. For basic parameters, let the combination of Fig. 1D be *d_b_, p_b_, q_b_*. The random geometric graph on **P**^∼^ was built with the same *d_b_* and with *p* = *p_b_* · *m*, with *m* = 0.04. The stochastic spread graph was then built with *q* = *q_b_* · (1 + *m*). Intuitively, this is to yield a small fraction (*m*) of inter-regional edges on top of the local connectivity. We found that, as expected, the connection probability was largely identical to the graph with only local connectivity below 500*µm*, but beyond that it flattened out, indicating the presence of long-range connections (Fig. 4B). These were also clearly visible, with topographical mapping in the two-dimensional connection probabilities (Fig. 4C). Next, we analyzed the simplicial structure of the two graphs and how the addition of inter-regional connections affected it. We found the overall number of simplices to be slightly lower with long-range connections in place (Fig. 4D). However, the simplices were much more likely to contain neurons from both sides of the region divide when long-range connectivity was present (Fig. 4E). In fact, large simplices were almost completely limited to only one side without it. As simplices have been shown to be motifs with functional implications [3, 24, 25], this demonstrates that the addition of long-range connectivity in such a model enhances inter-regional interactions.

## 4 Discussion

We have defined a type of graph that it constructed on top of the structure of a separate, underlying graph and used it to model the local connectivity of cortical circuitry. We have validated it by demonstrating that it recreates various features of connectivity that have been previously characterized in biological neuronal networks and found to be potentially functionally impactful. The features ranged from comparatively simple - e.g., distance-dependence of connectivity - to others that are difficult to obtain in random graph models - e.g., overexpression of directed simplices. We have further demonstrated that this type of graph can take on very different values for salient graph measurements, it is therefore flexible and can be used to model different types of circuitry. Notably, at the extreme ends it turns into an empty graph (stochastic spread on an empty graph) or an Erdos-Renyi graph (stochastic spread on a fully connected graph). We have demonstrated that after manual parameter optimization the type of graph can be used to model the network of synaptic connectivity of cortical circuitry, using an electron-microscopic reconstruction and a morphologically-detail model as references. Notably, the type of graph has longer tails for out-degree than in-degree distributions except in a portion of the parameter space with unreasonably high sparsity of connectivity (Fig. 1C). This prediction was confirmed in the data of the electron-microscopic MICrONS connectome (Figs. 2C2, S2C2).

On a conceptual level, the algorithm captures the growing and spreading of an axon through space, but with a spatial resolution given by the neuron density. As we were only interested in the wiring diagram, we would argue that higher resolution was not required. This insight greatly reduced the complexity of the implementation, allowing us to generate instances of a stochastic spread graph on 20,000 nodes in ∼1 second on a standard laptop computer.

When generating the stochastic spread graph, we used the idea that the probability to spread to a candidate, *r*, is determined by the number of candidates. An arguably more intuitive approach would use a fixed *r*. In that case, the spread would be subject to the rules of criticality, and we found that made it hard to parameterize and inflexible. It was also not needed for obtaining long-tailed degree distributions: At each step, the probability that the process terminates is the probability of obtaining 0 in a binomial distribution with *n*_binom_ equal to the number of candidates and *p*_binom_ = *q/n*_binom_. It is therefore a multiplicative process according to Piazza et al. [21].

As formulated, the algorithm emulates the spreading of the axon, but not the dendrite. This is based on our finding that the structure of the axon affects connectivity more than the dendrite. When repeating the analysis in Fig. 1B but for incoming instead of outgoing connectivity, we found a much weaker effect (Fig. S3).

As incoming connectivity is determined by the dendrites, this demonstrates that their effect the higher-order structure is not as strong as for axons. In the future, the algorithm can be extended to give dendrites a more prominent role as follows: One can represent larger dendritic morphologies by several points along its extent, e.g., along the apical dendrite. Each could represent, for example, one dendritic functional unit [28]. A random geometric graph is then built, but only the points representing somata are allowed to grow outgoing edges. This can be enforced by using a per-node bias of 0 for all others. In the final graph, the edges of a neuron would be the union of edges of all nodes associated with it. In models with spatially extended morphologies, this approach would also yield coarse-grained dendritic locations for synapses. The modification may also make the parameters enforcing the spatial structure of connectivity (*w_i_*, *w_o_*, *f_y_* and *w*_A_) superfluous, as larger dendritic morphologies would naturally have more inputs and from further away. Here, we wanted to present a solution for point neuron models, so the idea remains future work.

Finally, we have outlined a way to generalize the approach to model long-range inter-regional connectivity in addition to local connectivity. In our example, all neurons were allowed to grow long-range connections, which contradicts biological data [29]. This can be improved in the future by considering the edges of the random geometric graph for long-range connectivity (G(**P**^∼^)) only for projecting neuron classes. Additional future work will be finding improved ways of expressing **P**^∼^. In broad strokes, the strengths of voxelized or regional connectivity can serve as the basis of proximity in **P**^∼^. This could be calculated with existing dimensionality-reduction techniques. We note that the dimensionality of **P**^∼^ does not have to be 3. Furthermore, long-range connectivity may also be subject to additional hierarchical structure. One could experiment with building a stochastic spread graph on a stochastic spread graph or a stochastic geometric spread graph, i.e., 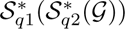 to capture this organization.

## Data and code availability

The code to generate SGSGs and to conduct the analyses of this manuscript is found in the following github repository: github.com/MWolfR/local_connectivity_model. The representation of the MICrONS connectome used as reference can be found on Zenodo under the following doi: 10.5281/zenodo.13849415. The representation of the connectivity of the model of Reimann et al. [14], also used as reference, can be found on Zenodo under the following doi: 10.5281/zenodo.10079406.

## Supplementary Material

**Fig. S1:**
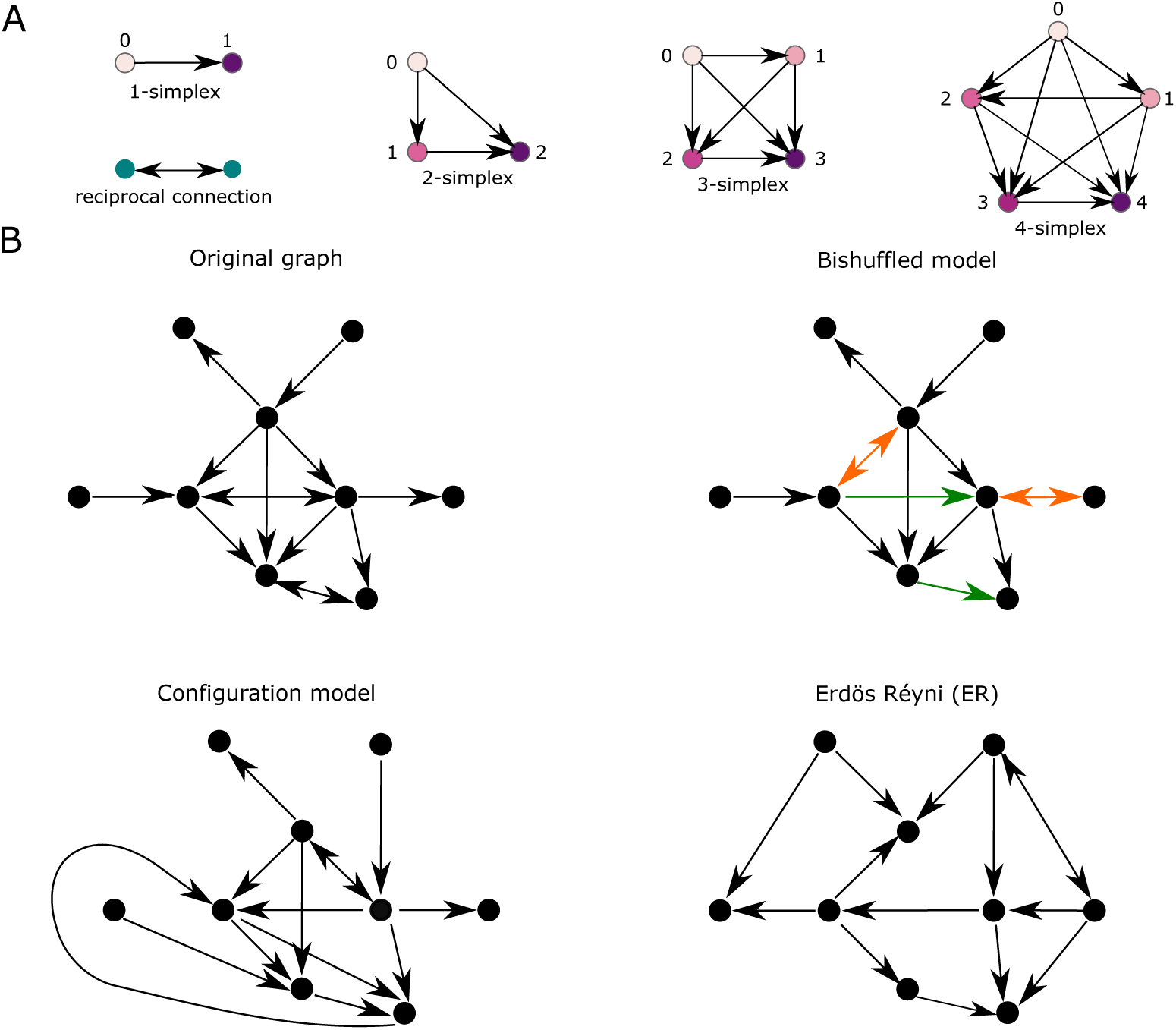
Simplex motifs and stochastic controls. **A** Simplex motifs for dimensions 1 through 5. These are fully connected feedforward motifs in which all nodes are connected such that there exists an ordering where there is an edge from *v* to *u* whenever *v < u*. In the diagram, this ordering is both labeled and reflected in the color scheme, transitioning from lighter to darker hues. **B** Different stochastic controls of an given original graph (top left). The *bishuffled* control alters only the positions of reciprocal (bidirectional) connections. Green edges indicate those that were reciprocal in the original, red edges indicate those that become reciprocal in the control. The *configuration model* and *Erdos–Rényi (ER)* models generate random graphs with the same number of edges as the original. The configuration model additionally preserves the in-degree and out-degree of each node, while ER scatters them at random across all edges.

**Fig. S2:**
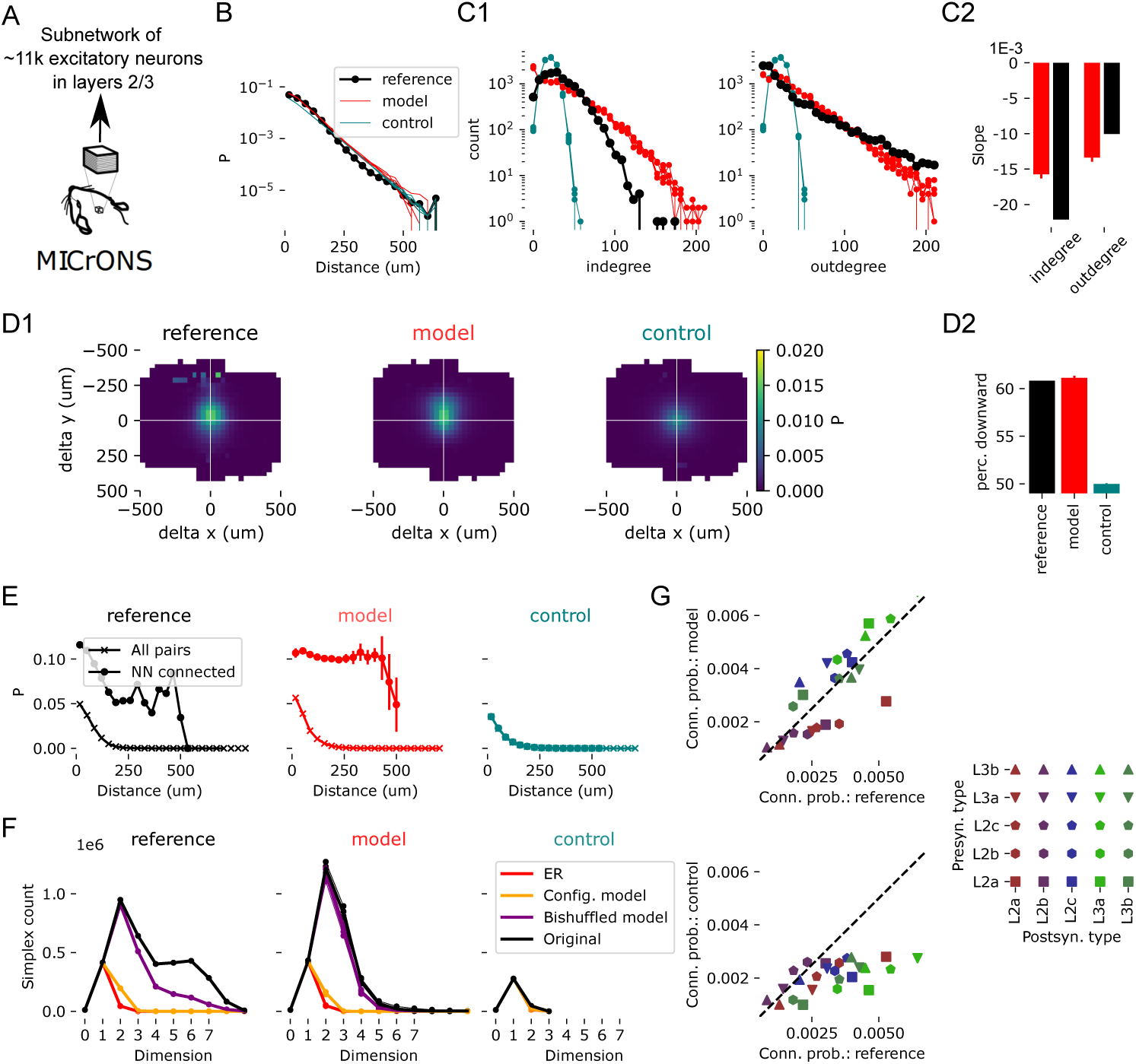
Comparison of a fitted SGSG* model to biological connectivity in L2/3. Everything as in Fig. 2, but using connectivity between the excitatory neurons in layers 2 and 3 of the MICrONS data as reference. Parameters *p*, *d* and *k* generalized from layers 4/5. *f_y_*, *q* and *w*_A_ re-fit to the data. Pearsonr for the data in panel F: 0.69 for reference vs. model, 0.42 for reference vs. control.

**Fig. S3:**
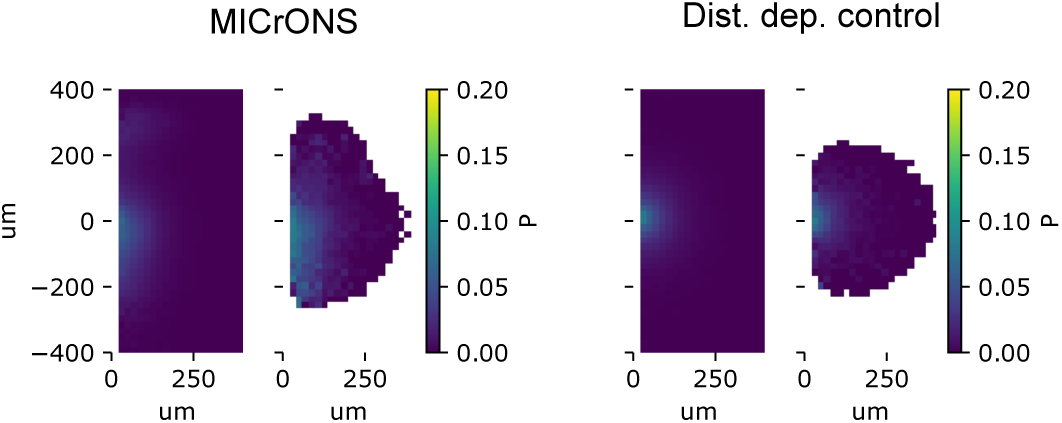
Nearest neighbor effect for incoming connectivity. Connection probability, conditional on the nearest neighbor being connected as in Fig. 1B, but for incoming connectivity instead of outgoing. Therefore, this analysis captures the impact of the physicality of dendrites instead of axons.

## Notes

https://github.com/MWolfR/local_connectivity_model

### Competing Interest Statement

The authors have declared no competing interest.

